# INSULIN-LIKE GROWTH FACTOR I SENSITIZATION REJUVENATES SLEEP PATTERNS IN OLD MICE

**DOI:** 10.1101/2021.02.23.432494

**Authors:** J.A. Zegarra-Valdivia, J. Fernandes, A. Trueba-Saiz, M.E. Fernandez de Sevilla, J. Pignatelli, K. Suda, L. Martinez-Rachadell, A.M. Fernandez, J. Esparza, M. Vega, A. Nuñez, I. Torres Aleman

## Abstract

Sleep disturbances are common during aging. Compared to young animals, old mice show altered sleep structure, with changes in both slow and fast electrocorticographic (ECoG) activity and fewer transitions between sleep and wake stages. Insulin-like growth factor I (IGF-I), which is involved in adaptive changes during aging, was previously shown to increase ECoG activity in young mice and monkeys. Furthermore, IGF-I shapes sleep architecture by modulating the activity of mouse orexin neurons in the lateral hypothalamus (LH). We now report that both ECoG stimulation and activation of orexin neurons by systemic IGF-I is abrogated in old mice. Moreover, stimulation of orthodromically activated LH neurons by either systemic or local IGF-I in young mice is absent in old mice. As orexin neurons of old mice show markedly increased IGF-I receptor (IGF-IR) levels, suggesting loss of sensitivity to IGF-I, we treated old mice with AIK3a305, a novel IGF-IR sensitizer, and observed restored responses to IGF-I and rejuvenation of sleep patterns. Thus, disturbed sleep structure in aging mice may be related to impaired IGF-I signaling onto orexin neurons, reflecting a broader loss of IGF-I activity in the aged mouse brain.

## Introduction

Sleep disturbances are so common during aging that whether they are an inherent component of the aging process is under debate ^1^. Importantly, sleep alterations present together with metabolic and cognitive impairments frequently found in aged individuals, but their contribution to pathology is unclear ^2,3^. Therefore, knowledge of the mechanisms underlying these age-associated changes in sleep architecture is of great interest.

We recently documented that circulating IGF-I, a hormone that participates in the aging process all along phylogeny ^4^, is also involved in the sleep/wake circadian cycle, as mice with reduced serum IGF-I present altered circadian electrocorticographic (ECoG) activity, among many other disturbances ^5^. Moreover, IGF-I directly shapes sleep architecture by modulating the activity of hypothalamic orexin neurons ^6^, a group of neurons in the lateral hypothalamus (LH) involved in sleep regulation ^7^. Significantly, aging is associated with reduced serum IGF-I levels in all mammalian species studied ^8^. However, whether IGF-I plays a detrimental or protective role in the aging brain is under debate ^9,10^. At any rate, loss of IGF-I activity in the aging brain has been previously documented by us ^11^ and many others ^12–14^. We hypothesized that orexin neurons in the aged brain might also lose sensitivity to this hormone, which, coupled with age-associated reduction of serum IGF-I would aggravate IGF-I loss-of-function, and eventually affect sleep/wake patterns.

We observed that systemic injection of IGF-I to old mice resulted in reduced c-fos expression in orexin neurons compared to young mice. Also, the responses of LH neurons to either systemic or local IGF-I were abrogated in old mice, which also points to loss of sensitivity to this growth factor. Since orexin neurons in old mice express higher IGF-IR levels, suggesting a compensatory mechanism to cope with lower brain IGF-I activity, we treated old mice with an IGF-IR sensitizer and found that responses to IGF-I recovered and sleep patterns rejuvenated. Thus, reduced IGF-I input to orexin neurons during aging contributes to age-associated sleep disturbances.

## Results

### Changes in sleep architecture in old mice

We first confirmed that changes in sleep structure during normal aging documented in humans ^15^ are also present in aged mice ^16^. Using ECoG recordings during the light period (corresponding to the inactive phase in mice, Figure 1A), we observed that old mice (>18 months old) have markedly different patterns of ECoG activity in both slow and high-frequency bands, as compared to young ones (<6 months old). Thus, the typical age-related decrease in δ activity ^15^, together with increases in θ, α, and β bands were observed (Figure 1B-E). The γ band was significantly depressed, specifically at Zeitgeber time (ZT) 19, suggesting that young animals woke up while old ones stayed sleeping (Figure 1F). ECoG analysis during the inactive phase shows a total mean decrease of δ activity (Figure 1G).

**Figure 1:**
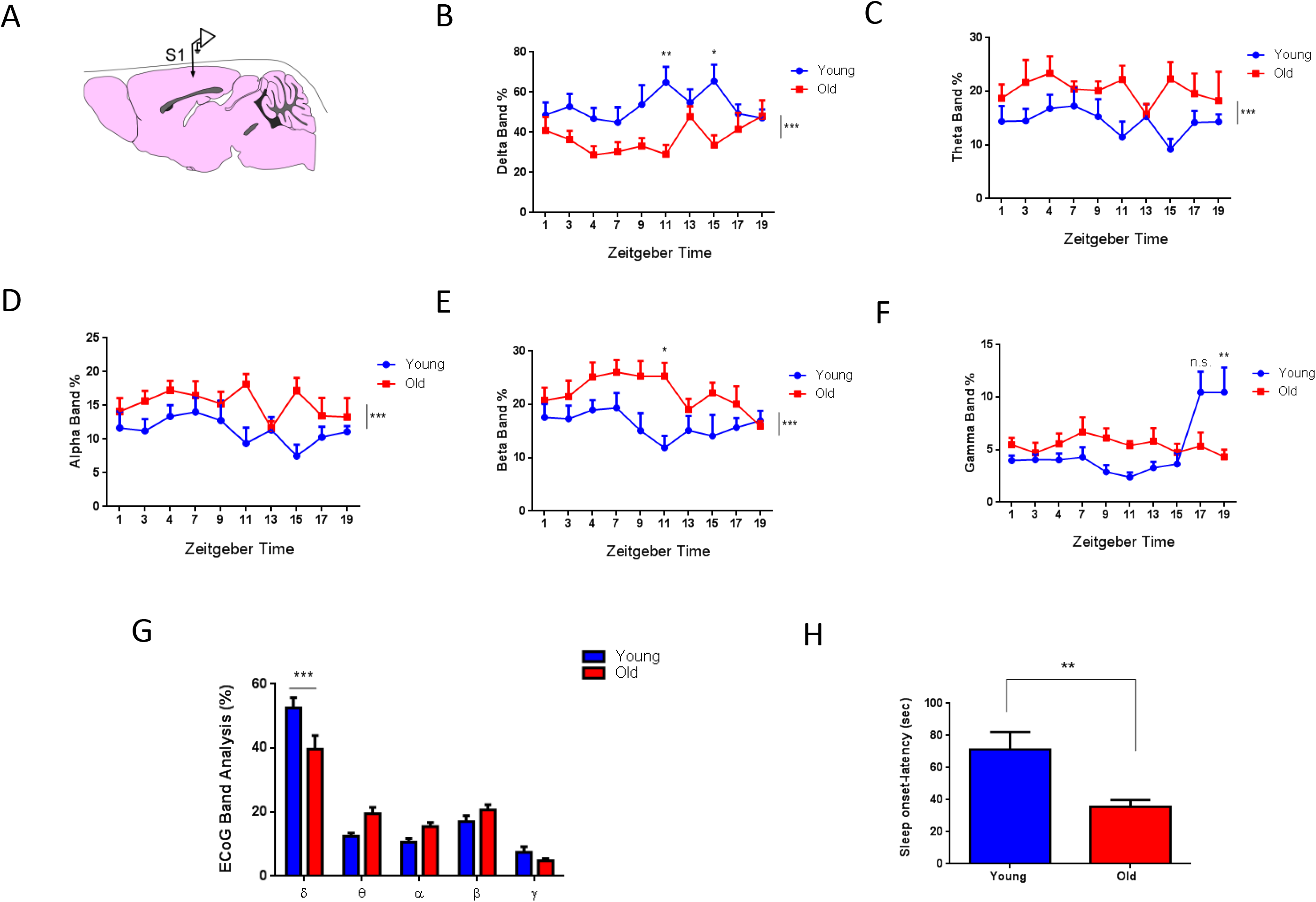
Altered sleep structure in old mice. **A,** Diagram of the intracranial localization of the electrodes in S1cortex; left hemisphere has the reference electrode in all cases. **B-F,** Sleep architecture during the light and dark period (ZT 1-19) determined by δ (C), θ (D), α (E), β (F), and γ (G) activity patterns in old (red line) and young mice (blue line). Old mice display lower delta activity, as compared to young ones, and an increase in fast-wave activity (young=8, old=4; male mice only, Two-Way ANOVA, and Sidak’s multiple comparison test; *p<0.05, **p<0.01, ***p<0.001). **G,** Average changes in ECoG bands during the passive phase (ZT 9 – 11) display significant differences in δ band (***p<0.0004, young=13, old=8, sex balanced, Two-Way ANOVA, and Sidak’s multiple comparison test). **H,** latency to sleep-onset was markedly reduced in old mice (**p<0.0099, young=11, old=8, sex balanced, Unpaired t-test, and Welch’s correction).

Next, we analyzed sleep onset latency and found that old mice show a significantly shorter latency to sleep (Figure 1H). We combined both sexes for these experiments, as ECoG patterns were similar in males and females (Suppl Figure 1A, B).

### Old mice show reduced sensibility to IGF-I

We previously reported that systemic administration of IGF-I (1 μg/g, ip) to young mice results in enhanced ECoG activity ^17^. ECoG responses to IGF-I induced a decrease of δ waves, while faster frequencies were significantly enhanced (Figure 2A). In contrast, old mice showed a slight increase of faster frequencies after ip IGF-I injection, reaching statistical significance only in the β frequency band (compare Figure 2A,B with Figure 2 in ref 17).

**Figure 2:**
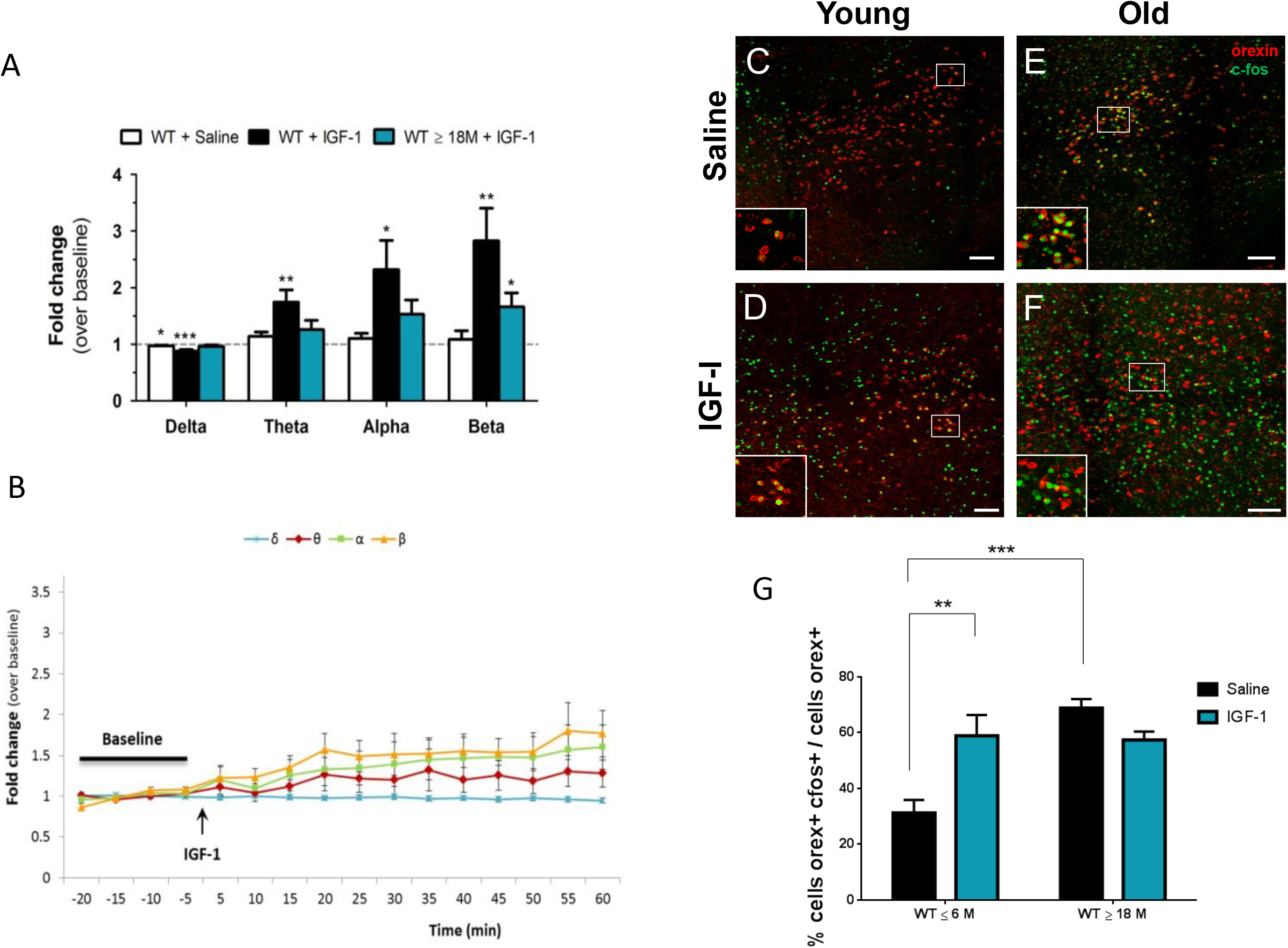
Loss of sensitivity to systemic IGF-I in aged mice. **A,** Average changes in ECoG bands from 20 to 60 min after administering IGF-I (1 μg/kg, ip) were compared with average baseline changes. ECoG response to IGF-I is lost in old mice (n = 7) in all bands, particularly in α, δ and θ compared to young mice (n=10). *p<0.05; **p<0.01; ***p<0.001, One-Way ANOVA). **B,** IGF-I administered to anesthetized old mice produced an attenuated increase in α, β and θ band frequencies in the ECoG (compare to Figure 2 in Trueba et al 2013). **C-F,** Representative photomicrographs of c-fos^+^/orexin^+^ cells immunostainings in young and old mice under saline and IGF-I condition. **G,** While young mice show increased c-fos expression in orexin neurons after ip IGF-I (n=6, **p<0.01), old mice show a not significant decreased expression (n=6, p=0.4948). Note that old mice (n=6) treated with saline show higher activation of c-fos than young saline mice (n=6, ***p<0.001; sex balanced, Two-Way ANOVA, and Sidak’s multiple comparison test).

Orexin neurons are a discrete cell population in the lateral hypothalamus that participate in sleep/wake regulation ^7,18^. Since IGF-I shapes sleep architecture through them ^6^, we speculated that these hypothalamic neurons might be involved in age-associated sleep disturbances by losing sensitivity to IGF-I. After systemic IGF-I injection, young mice responded with increased c-fos expression in orexin neurons, Conversely, whereas old mice had significantly higher basal c-fos immunoreactivity in orexin neurons than young mice, they did not respond to IGF-I (Figure 2C-G).

To elucidate whether reduced responses to systemic IGF-I are due to locally altered IGF-I signaling or reduced entrance of IGF-I from the periphery ^19,20^, we recorded responses to IGF-I in the PeF area of the lateral hypothalamus (LH) after either peripheral or local administration in both young and old mice. First, we activated LH neurons through the locus coeruleus (LC) as described before ^6^, and then we administered IGF-I either ip or locally (Figure 3A). While orthodromic stimulation of LH neurons elicited comparable evoked potentials and similar latency in young and old mice (Suppl Fig 1 C, D), this was not the case for their responses to IGF-I. Either after local (Figure 3B) or systemic (ip) IGF-I administration (Figure 3C), LH neurons of old mice responded with slightly reduced activity, whereas young mice responded with increased activity.

**Figure 3:**
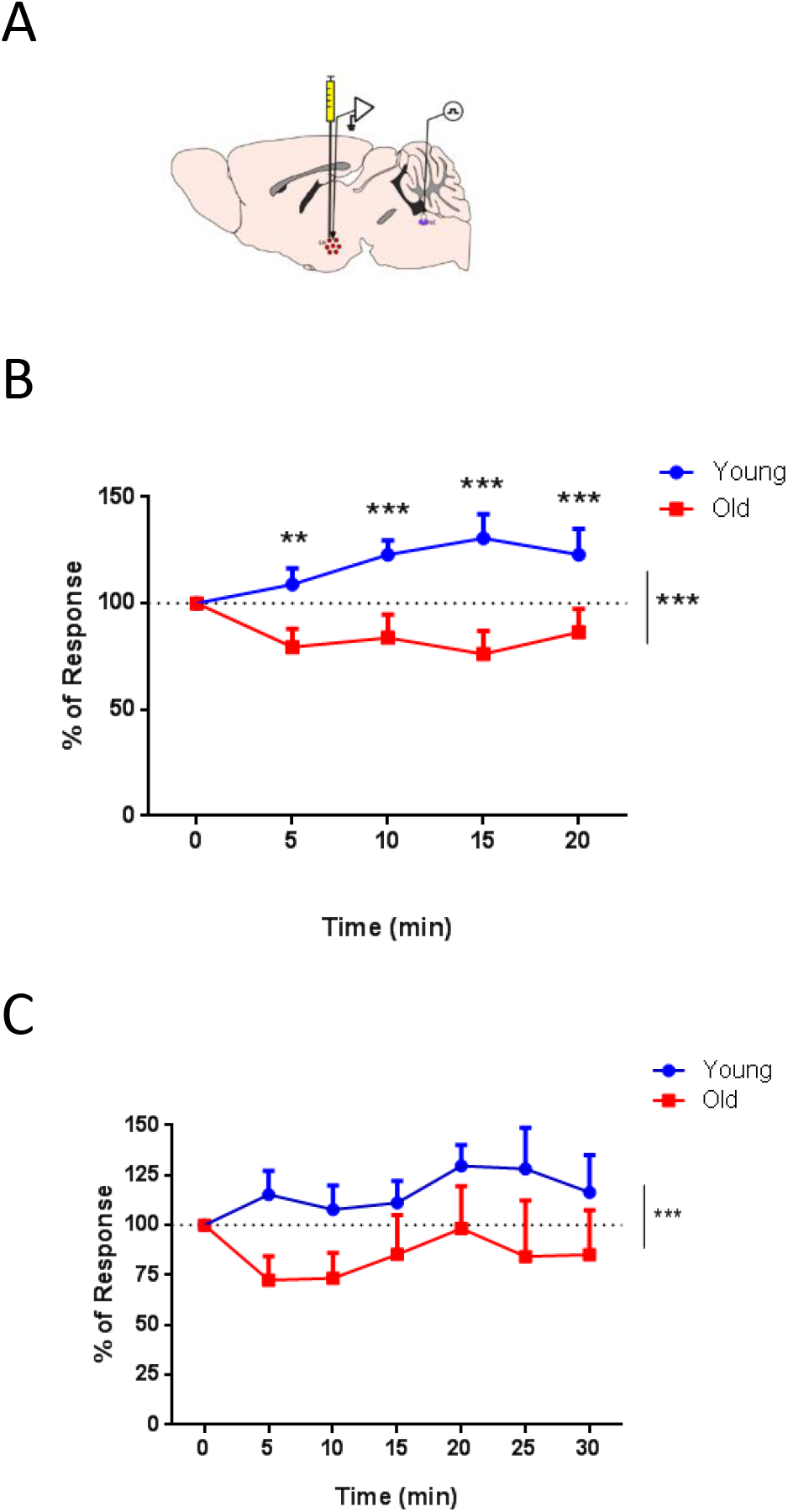
Orexin neurons in the lateral hypothalamus lose sensitivity to IGF-I in old mice. **A**, Diagram of the experimental design. A stimulating electrode was placed in the LC and a recording electrode in the PeF at the lateral hypothalamus (LH). **B**, Young, but not old mice responded to local IGF-I application in the PeF (arrow, 10 μM, 0.1 μl) after LC stimulation. Time course showing the evoked potential of orthodromic impulses after electrical stimulation of the LC in both experimental groups expressed as a percentage of basal responses at time 0 (at 5 min **p< 0.0053; at 10 min ***p<0.001, 15 min ***p<0.001, and 20 min ***p<0.001; n=13 per group; sex balanced, Two-Way Repeated Measure ANOVA, Sidak’s Multiple comparison test). **C,** Intraperitoneal injection of IGF-I (arrow, 1μg/g) increased neuronal activity in PeF orexin neurons after LC stimulation in young but not old mice (***p=0.0003; young=9; old=7; sex balanced, Ordinary Two-Way ANOVA, Sidak’s Multiple comparison test).

### IGF-IR sensitization recovers orexin responses to IGF-I

Altered responses to local administration of IGF-I suggests changes in orexin sensitivity to this growth factor. Thus, we examined IGF-IR expression in these neurons using double immunocytochemistry for orexin and IGF-IR (Figure 4A). Image analysis revealed a marked increase in double labelled IGF-IR/orexin cells in old mice (Figure 4B), while the number of orexin neurons (mean per field) was similar in both age groups (Figure 4C).

**Figure 4:**
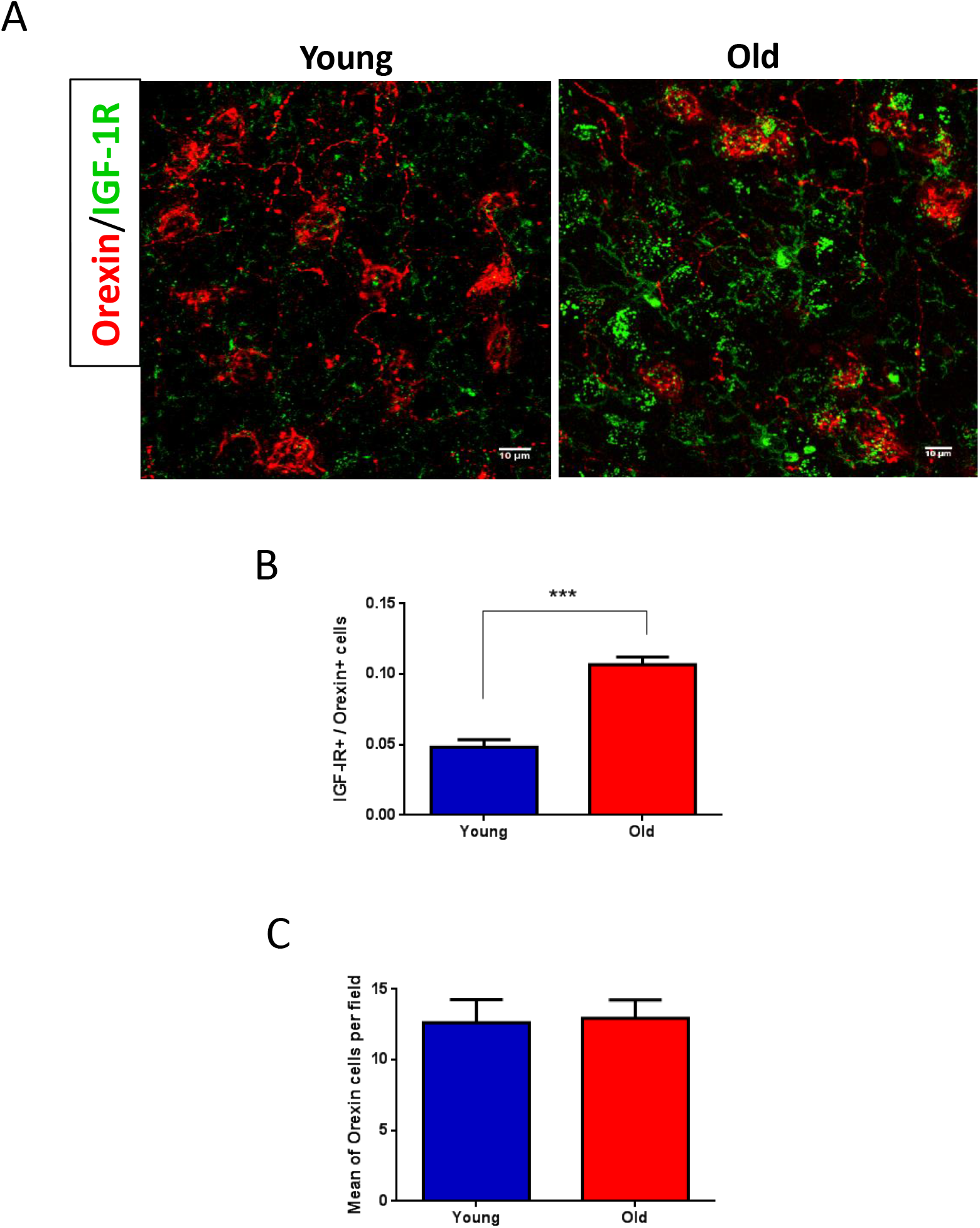
Increased expression of IGF-IR in orexin neurons of old mice. **A,** Representative photomicrograph of the mouse hypothalamus showing staining of orexin (red) and IGF-IR (green) in the PeF area of the lateral hypothalamus (LH). **B,** Co-localization ratio of IGF-1R/orexin cells in young and old mice (***p=0.0001, young=32, old=26; male-only, Unpaired t-test, and Welch’s correction). **C,** Mean orexin immunoreactivity per field in young and old mice, as detected by immunocytochemistry (young=32, old=26; p=0.8766, Unpaired t-test).

To confirm a loss of sensitivity to IGF-I in the old brain and at the same time assess a potential treatment for age-associated sleep disturbances, we treated aging mice with AIK3a305 (20 mg/kg/day; ip, 1 month), a novel IGF-IR sensitizer (Suppl Fig 2A) that crosses the blood-brain-barrier (Suppl Fig 2B) and found that c-fos responses to IGF-I were recovered to youthful levels (Figure 5A, B).

**Figure 5:**
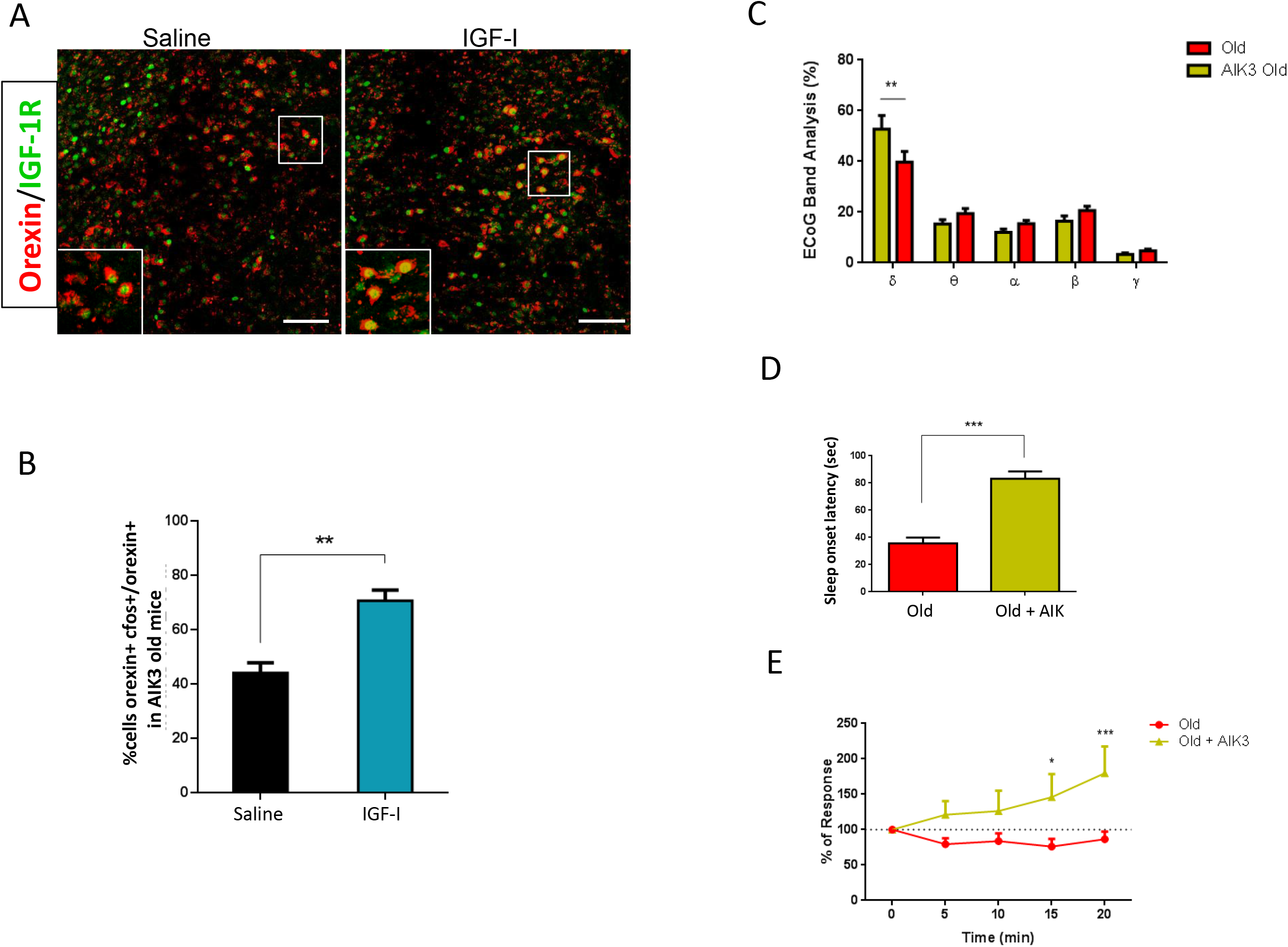
IGF-IR sensitization with Aik3a305 recovers orexin responses to IGF-I and rejuvenates ECoG patterns in old mice. **A,** Representative photomicrograph of orexin^+^ (red) and c-fos^+^ (green) cells under saline and IGF-I condition in old AIK3-treated mice. Scale bars: 100 μm. **B,** Counting of double-stained c-fos^+^/orexin^+^ cells after AIK3 treatment of old mice shows a recovery of the c-fos response to IGF-I (n=3 per group, **p=0.0082, sex balanced, Unpaired t-test, and Welch’s correction). **C,** Average ECoG bands during the passive phase (ZT 9 – 11) display significant differences in the δ band in old AIK3-treated mice, compared to old vehicle-injected mice (**p=0.0085, old AIK3=14, old=8, sex balanced, Two-Way ANOVA, and Sidak’s multiple comparison test). **D,** Treatment with AIK3 normalized sleep-onset latency (***p<0.001, n=7 per group, Unpaired t-test, and Welch’s correction). **E,** Old AIK3 mice recover responses to IGF-I in the LH after local application (10 μM, 0.1 μl) and LC stimulation. Time course showing the evoked potential of orthodromic impulses after electrical stimulation of the LC in both experimental groups expressed as a percentage of basal responses at time 0 (at 15 min *p=0.0109, and 20 min ***p=0.0004; n=13 per group; sex balanced, Two-Way Repeated Measure ANOVA, Sidak’s Multiple comparison test).

One month of treatment with AIK3a305 increased the percentage of δ waves during the inactive phase in old mice to levels seen in young animals (Figure 5C), while during the active phase, no differences were seen between groups (Suppl Fig 2C). The sleep-onset latency was also recovered by AIK3a305 treatment (Figure 5D). Further, the sensibility of orthodromically stimulated LH neurons to local IGF-I injection was also recovered (Figure 5E). Orthodromic evoked potentials showed no significant differences under basal conditions between the three groups (Suppl Figure 2D).

## Discussion

The present observations reinforce previous ones indicating that during aging, the brain loses sensitivity to IGF-I and suggest that this loss includes orexin neurons in the lateral hypothalamus. Since these neurons are involved in the sleep/wake cycle, and IGF-I influences sleep architecture through them ^6^, we may conclude that sleep activity in old mice is disturbed at least in part because IGF-I input to orexin neurons is compromised. Indeed, the presence of higher IGF-IR levels in aging orexin neurons suggests a compensatory mechanism to impaired IGF-I signaling due to resistance, deficiency, or both ^21^. This explanation is reinforced by the therapeutic actions of AIK3a305, a novel sensitizer of IGF-IR. The combined loss of sensitivity to IGF-I in orexin neurons and elsewhere, together with low serum IGF-I levels, will produce an overall reduction of IGF-I effects on the aged brain, as reflected by reduced ECoG responses to this growth factor.

Intriguingly, higher IGF-IR levels in the mouse brain have been associated with reduced longevity ^22^. Although it may seem counterintuitive, higher IGF-IR levels may point to a less efficient IGF-I system in the brain, like this and previous ^23^ observations suggest. Thus, higher expression of IGF-IR probably reflects post-receptor disturbances in IGF-I signal transmission ^24^, resulting in reduced activation of the pathway. Since IGF-I downregulates the expression of its own receptor in the brain ^25^, a rebound increased IGF-IR synthesis seems to take place. Other age-associated changes leading to aberrant IGF-IR function may also be involved, as reported for the brain insulin receptor ^26^.

The precise role of IGF-I in the aging process is controversial, albeit mainstream thinking supports the notion that its function, together with other members of the insulin family, is detrimental ^27–31^. While many observations of the beneficial actions of IGF-I indicate that this notion is not univocal ^32–36^, and is probably simplistic ^37,38^, in the particular case of brain aging, the situation is even more confusing, with both detrimental ^39,40^ and beneficial actions of IGF-I ^41–45^ profusely supported by data. Nevertheless, while these observations indicate that loss of IGF-I activity in the aging mouse brain disturbs sleep patterns, we cannot rule out potential beneficial effects of this loss in other aspects of brain function.

In summary, we observed a loss of sensitivity to IGF-I during brain aging, leading to dysregulated orexin function, which impacts on sleep patterns. Since IGF-IR sensitization recovered brain responses to IGF-I and rejuvenated sleep patterns, these results may provide new therapeutic avenues for age-related sleep disturbances.

## Material and Methods

### Materials

Antibodies used in this study include rabbit polyclonal anti-c-Fos (Abcam ab190289), rabbit polyclonal anti-IGF-I Receptor-β (Santa Cruz 713/AC) and rabbit anti-IGF-I receptor β XP (Cell Signaling Technology, USA), orexin polyclonal mouse antibody (Santa Cruz 80263), orexin polyclonal rabbit antibody (Abcam ab 6214), anti-phosphoSer Akt (Cell Signaling, 9271S), and monoclonal anti-phosphotyrosine (clone PY20, BD Transduction Laboratories, USA). Human recombinant IGF-I was from Pre-Protech (USA).

### Animals

Adult female and male young (4-5 months) and old (12-22 months) C57BL/6J mice (28-34g, Harlan Laboratories, Spain) were used. The estrous cycle of the female mice was not determined. Experiments were done during the light phase, except when indicated. Animals were housed in standard cages (48 × 26 cm^2^, 5 per cage) and kept in a room with controlled temperature (22°C) under a 12-12h light-dark cycle. Mice were fed with a pellet rodent diet and water *ad libitum*. Animal procedures followed European guidelines (2010/63, European Council Directives) and were approved by the local Bioethics Committee (Government of the Community of Madrid, Proex 112/16).

### Drug administration

IGF-I was dissolved in saline, and intraperitoneally (ip) injected (1μg/g body weight). In a subset of experiments, mice were processed for immunocytochemistry (see below) one or two hours after ip IGF-I injection to detect the expression of phospho-Akt or c-fos, respectively, whereas, in other experiments, PeF recordings were carried out immediately after ip injection. Alternatively, IGF-I was locally delivered in the PeF (10 nM; 0.1 μl; coordinates from Bregma: A, −1.95; L, 1 and depth, 4.0 - 4.5mm), and injected animals were then submitted to electrophysiological recordings, as described below. Doses were selected based on previous work using pharmacological injections for both systemic and local/intraparenchymal administration. AIK3a305 (Allinky Biopharma, Spain) or the vehicle (DMSO in saline) were injected intraperitoneally (ip) at a dose of 20 mg/kg/day for one month.

### Electrophysiological recordings

Mice were anesthetized with isofluorane (2% induction; 1–1.5% in oxygen, maintenance doses), placed in a David Kopf stereotaxic apparatus (Tujunga, CA, USA) in which surgical procedures and recordings were performed, with a warming pad (Gaymar T/Pump, USA) set at 37°C. Local anesthetic (lidocaine 1%) was applied to all skin incisions and pressure points. An incision was made exposing the skull, and small holes were drilled in the skull. Tungsten macroelectrodes (<1 MΩ World Precision Instruments, USA) were used to record the local field potential and the evoked potential in the PeF (coordinates from Bregma: A, −1.95; L, 1 and depth, 4.0 - 4.5mm).

Recordings were filtered (0.3–50 Hz) and amplified via an AC preamplifier (DAM80; World Precision Instruments). To elicit evoked potentials, the LC was stimulated using 120 μm diameter stainless steel bipolar electrodes (World Precision Instruments, coordinates from Bregma: A, −5.4; L, 1 and depth, 4.0 - 4.5mm). Electrical stimulation was carried out with single square pulses (0.3 ms duration and 20-50 μA intensity, delivered at 1 Hz; Cibertec stimulator, Spain). After a basal recording, IGF-I was injected either locally (young=13, old=13), or systemically (young= 9, old=7). Both sexes were included.

For recordings in freely moving animals, mice were anesthetized as indicated above and placed in a stereotaxic device. The skin was cut along the midline, and a craniotomy was made (0.5 mm diameter) in the primary somatosensory area (S1). A stainless-steel macro-electrode of <0.5 MΩ was placed without disrupting the meninges to register the cortical electrical activity (ECoG), using a DSI Implantable Telemetry device (Data Sciences International, USA). After surgery, mice remain in their cages for a minimum of 4 days to recover.

ECoG recordings were performed for 15 minutes from ZT1-ZT19, every 2 hours. Signals were stored in a computer using DSI software and filtered off-line between 0.3–50 Hz with Spike 2 software. ECoG segments of 5 minutes were analyzed using the Fast Fourier Transform algorithm to obtain the power spectra. The mean power density was calculated for five different frequency bands that constitute the global ECoG: delta (0.3–5 Hz), theta (5–8 Hz), alpha (8–12 Hz), beta (12–30 Hz), and gamma bands (30–50 Hz). The total power of the five frequency bands was considered 100%, and the percentage of each frequency band was calculated for the 15 minutes in each time point. We used eight young and four old mice to determine the ECoG profile during the active and passive phases.

To assess sleep onset-latency, segments of 30 sec of the ECoG recording were analyzed according to the presence of slow waves (0.3–5 Hz), fast waves (>12 Hz), and mouse movements. We measured the latency to sleep onset from the time the animal was placed on the platform until the absence of movement and the appearance of delta waves. ECoG recordings were performed at different time points through a remote computer.

### Data Analysis

Evoked potentials elicited by LC electrical stimulation (20-50 μA; 0.3 ms duration; at 1 Hz) were calculated. The peak latency was calculated as time elapsed between the stimulus onset and the peak of the second evoked potential wave (orthodromic response, with a latency of 3.5 ± 0.81 ms). To quantify the orthodromic response, the area under the positive wave curve was measured from the beginning of the positive slope. The unit activity plots show the percentage of variation over the basal period (5 mins). Outliers and recordings that did not elicit at least 70% of increment activity were removed from the analysis to minimize interference from non-LH neurons.

### Immunoassays

#### Immunocytochemistry

Animals were deeply anesthetized with pentobarbital (50 mg/kg) and perfused transcardially with 0.9% saline and then 4% paraformaldehyde in 0.1 M phosphate buffer, pH 7.4 (PB). Coronal 50-μm-thick brain sections were cut in a vibratome and collected in PB 0.1 N. Sections were incubated in permeabilization solution (PB 0.1N, 1% Triton X-100), followed by 48 hours incubation at 4°C with primary antibody (1:500) in blocking solution (PB 0.1N, 0.1% Triton X-100, 10% normal horse serum). After washing three times in 0.1 PB, Alexa-coupled mouse/rabbit polyclonal secondary antibodies (1:1000, Molecular Probes, USA) were used. Finally, a 1:1000 dilution in PB of Hoechst 33342 was added for 3 minutes. Slices were rinsed several times in PB, mounted with gerbatol mounting medium, and allowed to dry. The omission of the primary antibody was used as a control.

#### Western Blotting

Assays were performed as described in detail elsewhere ^46^. Densitometry analysis of blots was performed using the Odyssey system (Lycor Biosciences, USA). A representative blot is shown from a total of at least three independent experiments.

### Cell Image Analysis

Confocal analysis was performed in a Leica (Germany) microscope. For double-stained orexin/c-fos cell counting, four sections per animal were scored using the Imaris software, as described ^6^. Alternatively, for double labelled IGF-IR/orexin cells, confocal images were segmented via k-means clustering with the number of clusters set to 5 (*k* = 5) using MATLAB ^47^ to obtain binary masks for each channel dividing background from foreground. Red channel masks (Orexin antibody) were further processed by individualizing unconnected regions, putative neurons, and eliminating isolated components with areas smaller than 25 μm^2^. The number of neurons in each mask was manually assessed. Green channel masks (IGF-IR antibody) were also visually inspected to ensure proper segmentation. Red and green channels were superposed to study the area of co-localization. For each segmented neuron (red channel), the fraction of its area that co-localized with IGF-IR antibody (green channel) was measured as an indirect quantification of the amount of receptor present in that neuron. We referred to this fraction as the co-localization index. The co-localization index of each image was computed as the mean co-localization index of the neurons present in that image. Lastly, the final co-localization index for each condition was computed as the mean co-localization index of all the images belonging to that condition.

Images with only secondary antibody stainings were used to check specificity of the secondary antibody ^48^ and segmented as described above. Specificity was evaluated by measuring the segmented receptor’s area and the number of segmented neurons and comparing the results with tissue images that underwent a complete immunohistochemistry protocol.

### Astrocyte cultures

Astroglial cultures were prepared as described in detail elsewhere ^49^ from postnatal (day 1-2) brains. After the forebrain was removed and mechanically dissociated, the mixed cell suspension was centrifuged and plated in DMEM/F-12 (Life Technologies) with 10% fetal bovine serum (Life Technologies) and 100 mg/ml of antibiotic-antimycotic solution (Sigma-Aldrich, Spain). Cells were maintained for 2 weeks at 37°C, 5% CO_2_ and re-plated at 10^5^ cells/cm^2^ in a 12-multiwell plate and grown until 80% confluency. On the day of the experiment, cells were washed twice with warm PBS and medium without FCS was added. Then, LPS was added (1μg/ml) and cells incubated for 12h at 37°C. After LPS treatment, cells were washed twice with PBS, and a fresh medium without FCS was added. AIKa305 was added (27nM) in DMSO for 3 hours at 37°C. Controls received DMSO alone. After AIK treatment, IGF-I (1 nM) was added, and cells incubated for 1h at 37°C. Then, plates were placed on ice for 5 minutes, washed and 100 μl of Lysis buffer containing proteases and phosphates inhibitors (Merck) with Laemmli loading buffer 1X, was added and cells scrapped on ice. Samples were frozen at −20°C until use.

### Statistical Analysis

Statistical analysis was performed using GraphPad Prism 6 software (San Diego, CA, USA) and R Package (Vienna, Austria). Depending on the number of independent variables, normally distributed data (Kolmogorov-Smirnov normality test), and the experimental groups compared, we used either Student’s t-test, two-way ANOVAs, or Two-way repeated measure ANOVA, followed by Sidak’s multiple comparison test. For non-normally distributed data, we used the Mann-Whitney U test to compare two groups, Kruskal-Wallis or Friedman test, with Dunn’s multiple comparisons. As a post hoc analysis we used the Scheirer-Ray Test, a non-parametric alternative to multifactorial ANOVA. The sample size for each experiment was chosen based on previous experience and aimed to detect at least a p<0.05 in the different tests applied, considering a reduced use of animals. Results are shown as mean ± standard error (SEM) and *p* values coded as follows: *p< 0.05, **p< 0.01, ***p< 0.001. Animals were included in each experimental group randomly by the researcher.

## Acknowledgments

This work was funded by a grant from Ciberned and SAF2016-76462 (AEI/FEDER; MINECO). J.A. Zegarra-Valdivia acknowledges the financial support of the National Council of Science, Technology and Technological Innovation (CONCYTEC, Perú) through the National Fund for Scientific and Technological Development (FONDECYT, Perú). J. Fernandes received a post-doc fellowship from Fundação de Amparo à Pesquisa do Estado de São Paulo (FAPESP: # 2017/14742-0; # 2019/03368-5). Kentaro Suda is supported by Kobe University. We are thankful to M. Garcia for technical support.

## Author Contributions

JAZV conducted experiments, prepared figures, results and wrote part of the manuscript. JF, ATS, MEFS, JP, AMF conducted experiments and prepared figures. KS and LMR conducted experiments, MN provided experimental expertise, JE analyzed data, MV provided reagents and data on AIK3a305, AN designed and conducted experiments and wrote part of the manuscript, ITA designed the study and wrote the manuscript.

---

**Supplementary Figure 1: A,** No differences were seen in the ECoG band analysis of young male and female mice (≤ 6 months old). **B,** No significant differences were seen in the ECoG band analysis of old male and female mice (≥ 18 months old). **C,** No differences were seen in the latency of the evoked potential registered in the PeF of young and old mice after stimulation of the LC. **D,** No differences between groups in the basal evoked potential in LH after LC stimulation (One-way ANOVA, p=0.1028).

**Supplementary Figure 2: A,** AIK3 sensitizes astrocytes to IGF-I. After treatment with LPS (1 μg/ml), astrocytes become unresponsive to IGF-I, as determined by lack of phosphorylated Akt (pAkt) after 0.1 or 1 nM IGF-I. Astrocytes simultaneously receiving AIK3a305 (27 mM) regain sensitivity to IGF-I, as phosphorylated Akt was readily detected. A representative blot is shown. **B,** Concentration along time in the blood (blue trace) and brain (red trace) after oral administration of AIK3a305 (10 mg/kg) in adult male mice (Swiss albino). **C,** No differences were seen in the ECoG band analysis of young mice (≤ 6 months old), old mice (≥ 18 months old), and old mice treated with AIK3. **D,** No differences between groups in the basal evoked potential in LH after LC.

