## Supplementary Figures for "INSULIN-LIKE GROWTH FACTOR I SENSITIZATION REJUVENATES SLEEP PATTERNS IN OLD MICE"

### Slide 1
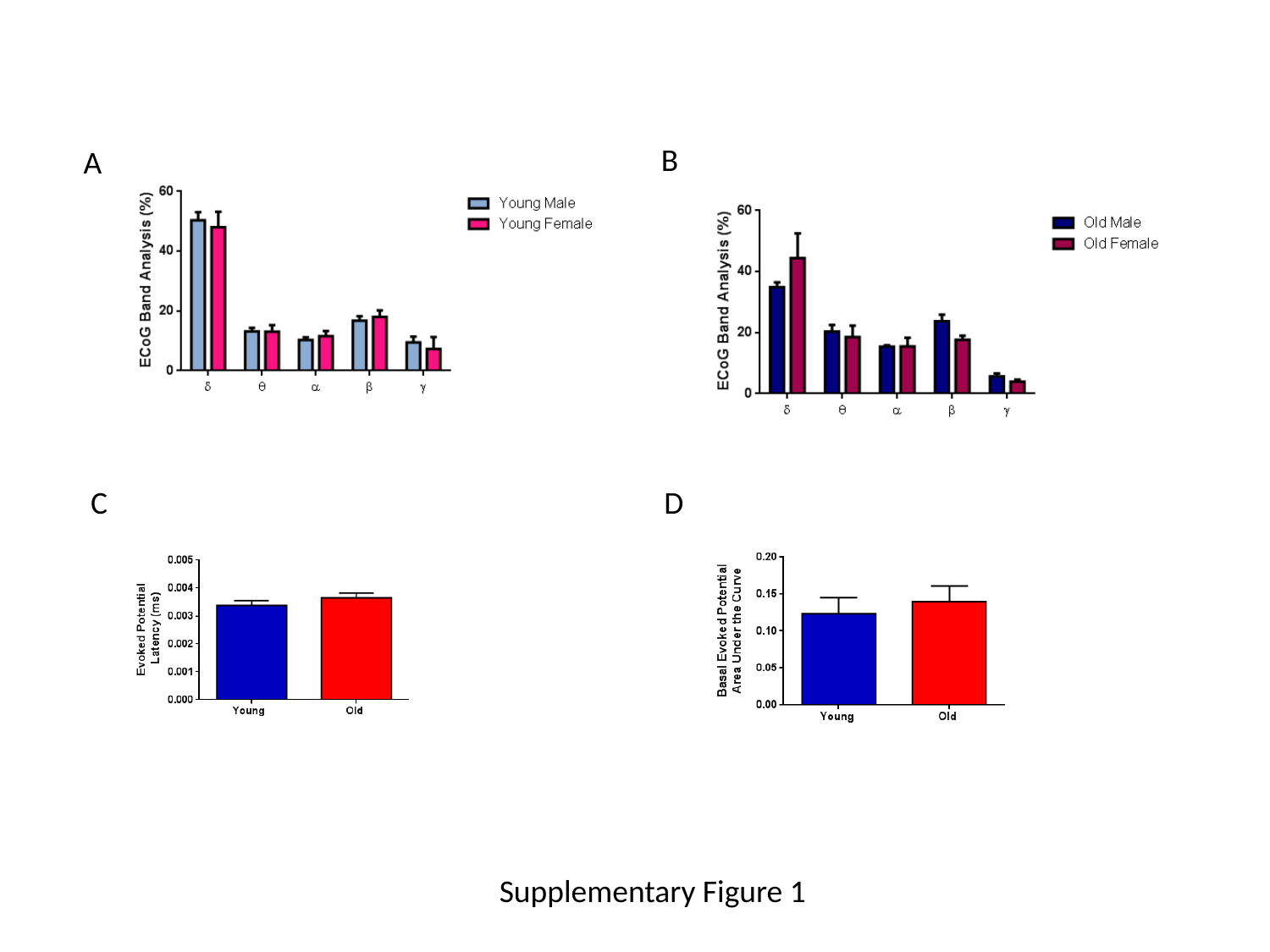

B
A
C
D
Supplementary Figure 1

### Slide 2
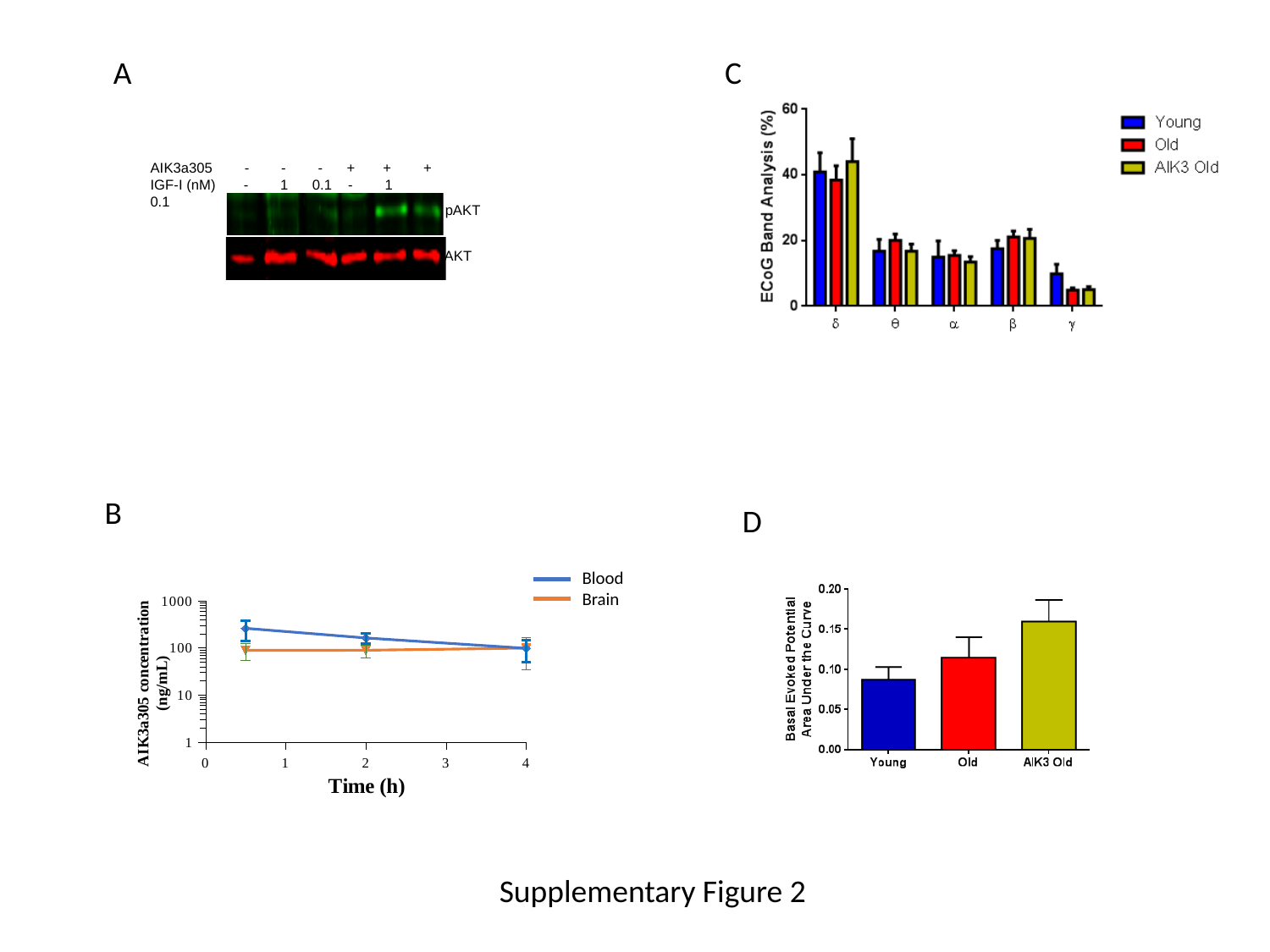

A
C
AIK3a305 - - - + + +
IGF-I (nM) - 1 0.1 - 1 0.1
pAKT
AKT
B
D
Blood
Brain
#### Chart
| Category | Plasma concentration (ng/mL) of ALK3a305 following Per oral administration to Swiss Albino mice (P0-10.0 mg/kg) | Brain concentration (ng/g) of ALK3a305 following Per oral administration to Swiss Albino mice (P0-10.0 mg/kg) |
|---|---|---|
Supplementary Figure 2
